# *SUPERMAN* prevents stamen formation and promotes stem cell termination in the fourth whorl of the Arabidopsis flower

**DOI:** 10.1101/107722

**Authors:** Nathanaël Prunet, Weibing Yang, Pradeep Das, Elliot M. Meyerowitz, Thomas P. Jack

## Abstract

The molecular and genetic networks underlying the determination of floral organ identity are well studied, but much less is known about how the flower is partitioned into four developmentally distinct whorls. The *SUPERMAN* gene is required for proper specification of the boundary between stamens in whorl 3 and carpels in whorl 4, as *superman* mutants exhibit supernumerary stamens but usually lack carpels. However, it has remained unclear whether extra stamens in *superman* mutants originate from an organ identity change in whorl 4 or the overproliferation of whorl 3. Using live confocal imaging, we show that the extra stamens in *superman* mutants arise from cells in whorl 4, which change their fate from female to male, while floral stem cells proliferate longer, allowing for the production of additional stamens.

## Introduction

Most of the adult body of flowering plants is generated post-embryonically from apical meristems, which are pools of undifferentiated cells at the tips of stems and roots. Divisions in the shoot apical meristem (SAM) allow for the continuous production of lateral organs on the flanks of the stem: leaves during the vegetative phase, then floral meristems (FMs) after the SAM transitions to the reproductive phase. FMs in turn generate the floral organs that comprise the flower.

Partition of the organism into distinct tissues and organs is a fundamental process of development in both animals and plants, yet it relies on different mechanisms in each kingdom. In animals, tissue separation is determined by cell surface cues that influence the adhesive properties of cells and their ability to interact with each other^1^. Unlike animal cells, however, plant cells are surrounded and connected to their neighbors by contiguous cell walls that prevent them from migrating. As new organs form, they are separated from surrounding tissues by a boundary, which consists of a group of cells with restricted growth that act as a physical barrier separating two different developmental programs^2^.

Unlike the SAM, which gives rise to lateral organs one at a time, in an iterative, spiral pattern, the FM semi-synchronously produces 16 floral organs, with four different identities, in four adjacent whorls. Floral organ identity is determined by the combinatorial action of four classes of MADS-box transcription factors, which form distinct complexes in the four floral whorls^3,4^. For instance, a combination of APETALA3 and PISTILLATA (AP3 and PI, class B), together with AGAMOUS (AG, class C) and SEPALLATA3 (SEP3, class E) specifies stamens in whorl 3, while complexes composed solely of AG and SEP3 trigger carpel development in whorl 4. Targets of these MADS-box transcription factors have been extensively studied, and downstream regulatory networks have been partially deciphered^4^. However, the mechanisms that underlie the patterning of the FM, with the generation of four distinct types of organs in such a constrained space and time, remain poorly understood. In particular, how boundaries between the floral whorls are established is still unclear. Here, we analyze the role of *SUPERMAN* (*SUP*) in defining the boundary between whorl 3 stamens and whorl 4 carpels.

*SUP* encodes a transcriptional repressor with a C2H2 zinc-finger DNA-binding domain and an EAR repression domain^5–8^, and is expressed at the boundary between whorls 3 and 4 ^6,9^. *sup* mutant flowers have numerous extra stamens, while carpel tissue is usually reduced or missing^10,11^. This phenotype is associated with the expansion of *AP3* and *PI* expression closer to the center of the FM compared to the wild type^10^. Overall, floral organ number is higher in *sup* flowers than in the wild type, indicating an increase in cell proliferation in developing *sup* flower buds. Although *SUP* was first characterized a quarter century ago, there are still two conflicting models to explain *SUP* function and the developmental origin of the *sup* phenotype. Here, we refer to these two models as “whorl 3” and “whorl 4” models, based on the whorl where the extra stamens in *sup* mutant flowers hypothetically form. The whorl 4 model proposes that *SUP* functions to prevent ectopic expression of *AP3* and *PI* in whorl 4. According to this model, ectopic *AP3/PI* expression in whorl 4 of developing *sup* flowers triggers the formation of stamens instead of carpels, and prolongs cell proliferation in the FM^10,11^. Conversely, the whorl 3 model proposes that *SUP* controls the balance of cell proliferation between whorl 3 and 4, and suggests that production of extra stamens in *sup* mutant flowers results from increased cell proliferation in whorl 3 at the expense of whorl 4^6,12^. In this study, we used live confocal imaging to investigate the developmental basis of the *sup* phenotype. We show that extra stamens in *sup* mutant flowers arise from a subset of whorl 4 cells that switch identity from female to male, as predicted by the whorl 4 model.

## Results

### SUP is expressed on both sides of the boundary between whorl 3 and 4

We generated a *gSUP-3xVenusN7* translational *SUP* reporter that complements the *sup-1* mutant phenotype. The SUP protein is first detected at stage 3, in cells adjacent to the boundary between whorl 3 and 4, inside of lateral sepal primordia (Fig. 1A-B; stages as described in^13^), and quickly expands to form an oblong ring approximately 3-4-cell wide, and longer medially than laterally (Fig. 1, A and C). At early stage 5, SUP is detected on both sides of the boundary between whorls 3 and 4, which at this stage forms a groove between the developing stamen primordia and the center of the flower (Fig. 1D). By late stage 5, *SUP* expression becomes restricted to a narrower band of cells at the boundary (Fig. 1A). *gSUP-3xVenusN7* fluorescence appears to peak at stage 4, before decreasing in intensity during stage 5 and becoming undetectable by late stage 6 (Fig. 1A). Overall, the *SUP* expression pattern resembles that of *AP3*, but *SUP* appears to be expressed closer to the center of the flower (Fig. 1, compare A and E). To determine more precisely where *SUP* is expressed relative to the boundary between whorl 3 and 4, we monitored the expression of *SUP* and class B genes simultaneously, using the *gSUP-3xVenusN7* reporter together with a *gAP3-GFP* (Fig. 1F) or a *gPI-GFP* (Fig. S1) translational reporter, or a *pAP3-CFPN7* transcriptional reporter (Fig. S2). *SUP* expression initiates shortly after that of *AP3* at stage 3 (Fig. 1F), and the first cells to express *SUP* also express *AP3* (Fig. 1G), indicating that *SUP* is initially expressed in whorl 3. However, from late stage 3 on, we observed cells that express *SUP* but do not express *AP3* or *PI* (Fig. 1H-I; Fig. S1; Fig. S2), demonstrating that *SUP* expression expands into whorl 4. At stages 4-5, SUP is clearly found on both sides of the boundary between whorls 3 and 4 (Fig. 1I; Fig. S1, B-D; Fig. S2, A1-A3). *SUP* expression overlaps with *AP3/PI* in whorl 3 at the boundaries between stamen primordia (Fig. S1D Fig. S2, B, D and F1-F3; Movie S1), as well as in a narrow, 1-2-cell wide band on the adaxial side of stamen primordia (Fig. 1I; Fig. S1D; Fig. S2, C and E1-E3; Movie S1). *SUP* is also expressed without *AP3/PI* in another narrow, 1-2-cell wide band in the outer part of whorl 4 (Fig. 1I; Fig. S1D; Movie S1). Together, these data clearly show that, contrary to earlier interpretations^6^, *SUP* is expressed on both sides of the boundary between whorl 3 stamens and whorl 4 carpels, and is not confined solely to whorl 3. Double fluorescence *in situ* hybridization experiments for *SUP* and *AP3* confirmed that this is also the case at the mRNA level (Fig. S3). Indeed, *SUP* protein levels appear higher in whorl 4, where class B genes are not expressed (Fig. S2, G1 and H). Similarly, *AP3* expression appears stronger in whorl 3 cells that do not express *SUP* (Fig. S2, G2 and I).

**Figure 1.**
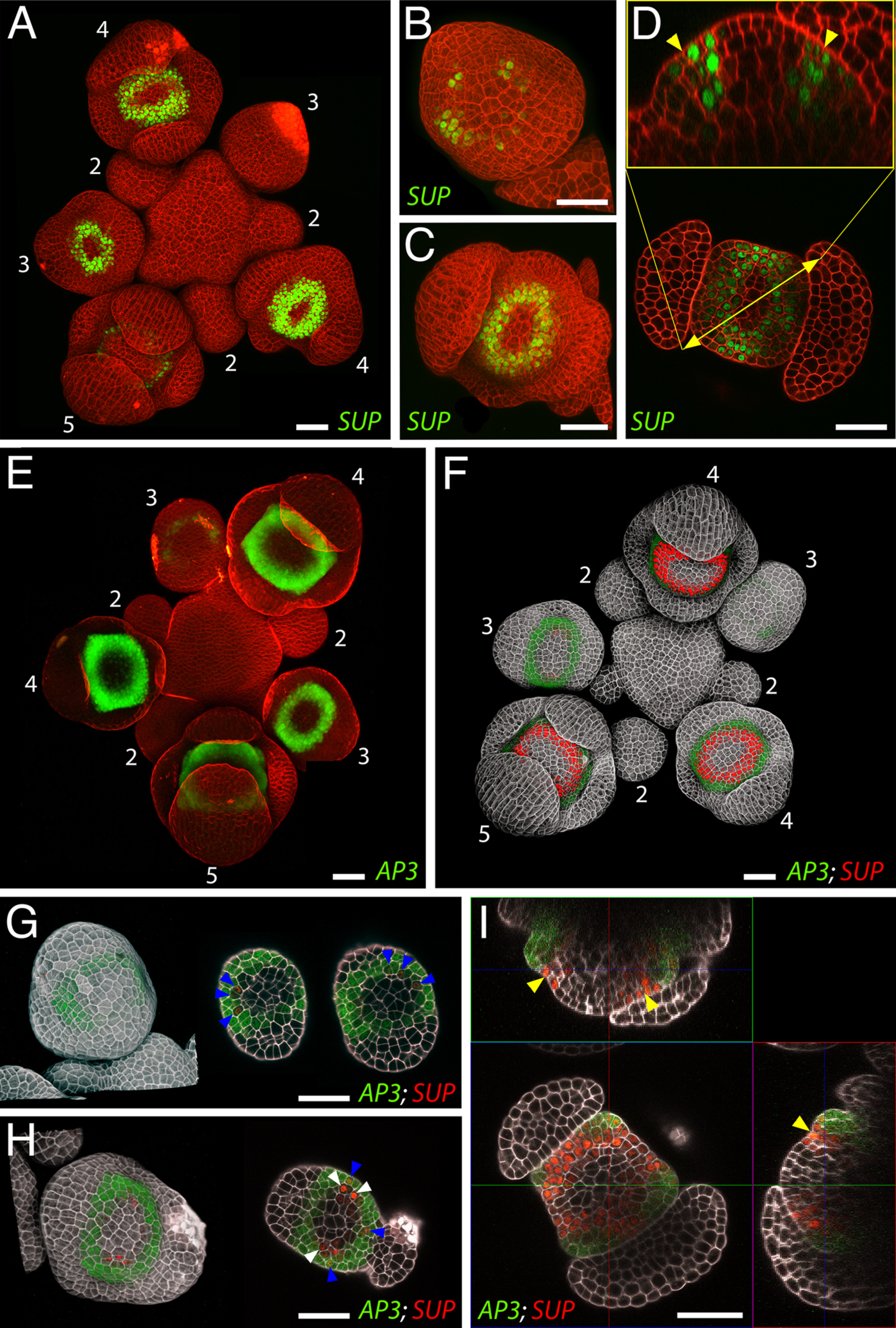
Expression of *SUP* and *AP3* in wild-type flowers. Expression of the *gSUP-3xVenusN7* (A-D) and *gAP3-GFP* (E) reporters separately (A-E), or together (F-I). (A, E and F) whole inflorescences; numbers indicate floral stages. (B-D and G-I) flower buds at early stage 3 (B and G), late stage 3 (H), stage 4 (C) and stage 5 (D and I). A, E, F and left panels in G and H show maximum intensity projections (MaxIPs). Lower panel in D, right panels in G and H and bottom left panel in I show slice views along horizontal planes. Upper panel in D, top left and bottom right panels in I show slice views along vertical planes. Yellow arrowheads indicate the position of the boundary between whorls 3 and 4, blue arrowheads mark cells that express both *AP3* and *SUP*, and white arrowheads mark cells that express *SUP* but not *AP3*. Bars = 25 µm.

To better understand where *SUP* is expressed relative to the positions where stamen and carpel primordia initiate, we examined plants expressing both the *gSUP-3xVenusN7* reporter and the *DORNROSCHEN-LIKE* (*DRNL*) *pDRNL-erGFP* reporter, which marks floral organ founder cells (Fig. S4, A-B)^14^. In particular, in whorl 3, *DRNL* is expressed in a ring that is reminiscent of *AP3* and *SUP* expression patterns (Fig. S4A), while in whorl 4, *DRNL* is expressed in two foci, which correspond to the sites of carpel initiation, as well as in two narrow arcs of cells connecting these foci, (Fig. S4, B and C)^14^. *SUP* and *DRNL* expression partially overlap in stamen primordia in whorl 3 (Fig. S4, A-B), while a narrow ring of *SUP* expression in whorl 4 directly surrounds *DRNL* expression in carpel founder cells in the center of whorl 4 (Fig. S4B).

### Extra stamens in sup-1 flowers arise from whorl 4 cells

To determine whether the extra stamens in *sup* mutant flowers arise from whorl 3 or whorl 4, we compared the expression of class B genes in wild-type and *sup-1* flowers using a *pAP3-3xVenusN7* transcriptional reporter (Fig. 2) and the *gAP3-GFP* (Fig. S5) and *gPI-GFP* translational reporters (Fig. S6). At stages 3-4, *AP3* expression appears similar in the wild type and in *sup-1* (Fig. 2, compare A and B; Fig. S5, compare A and B). However, by stage 5, both *AP3* and *PI* are expressed closer to the center of the flower in *sup-1* than in the wild type (Fig.2, compare A and B; Fig. S5, compare A and B; Fig. S6, compare A and B)^10^. While the fourth whorl of wild-type flowers shows no *AP3/PI* expression (Fig. 2, C and E; Fig. S6C), a narrow, 2-cell wide band of *AP3/PI* expression can be seen inside of the boundary between stamen primordia and the center of *sup-1* flowers at stage 5 (Fig. 2, D and F; Fig. S6D). At stage 6, the whole fourth whorl of wild-type flowers develops into carpel primordia (Fig. 2C)^13^. Conversely, in *sup-1* flowers, extra stamen primordia only start forming within the ring of extra *AP3*-expressing cells at stage 7, with a slight delay compared to wild-type carpels (Fig. 2G; stages for *sup-1* flowers were determined based on time elapsed after stage 5, which is the last stage at which wild-type and *sup-1* flowers are morphologically identical). As these extra stamens develop, *AP3* expression spreads again beyond the boundary of the primordia towards the center of *sup-1* flowers, forming another narrow ring of *AP3-*expressing cells, which later gives rise to more stamen primordia (Fig. 2K). This iterative process allows for the formation of several consecutive rings of stamens, sometimes resulting in flowers with more than 20 stamens. It is worth noting that *AP3* is never expressed throughout the center of *sup-1* flowers, which eventually develop into stunted, misshapen carpels or chimeric stamen/carpel organs^10,11^. Accordingly, organ primordia composed both of cells that express *AP3* and cells that do not, can often be seen in the center of developing *sup-1* flowers (Fig. S7).

**Figure 2.**
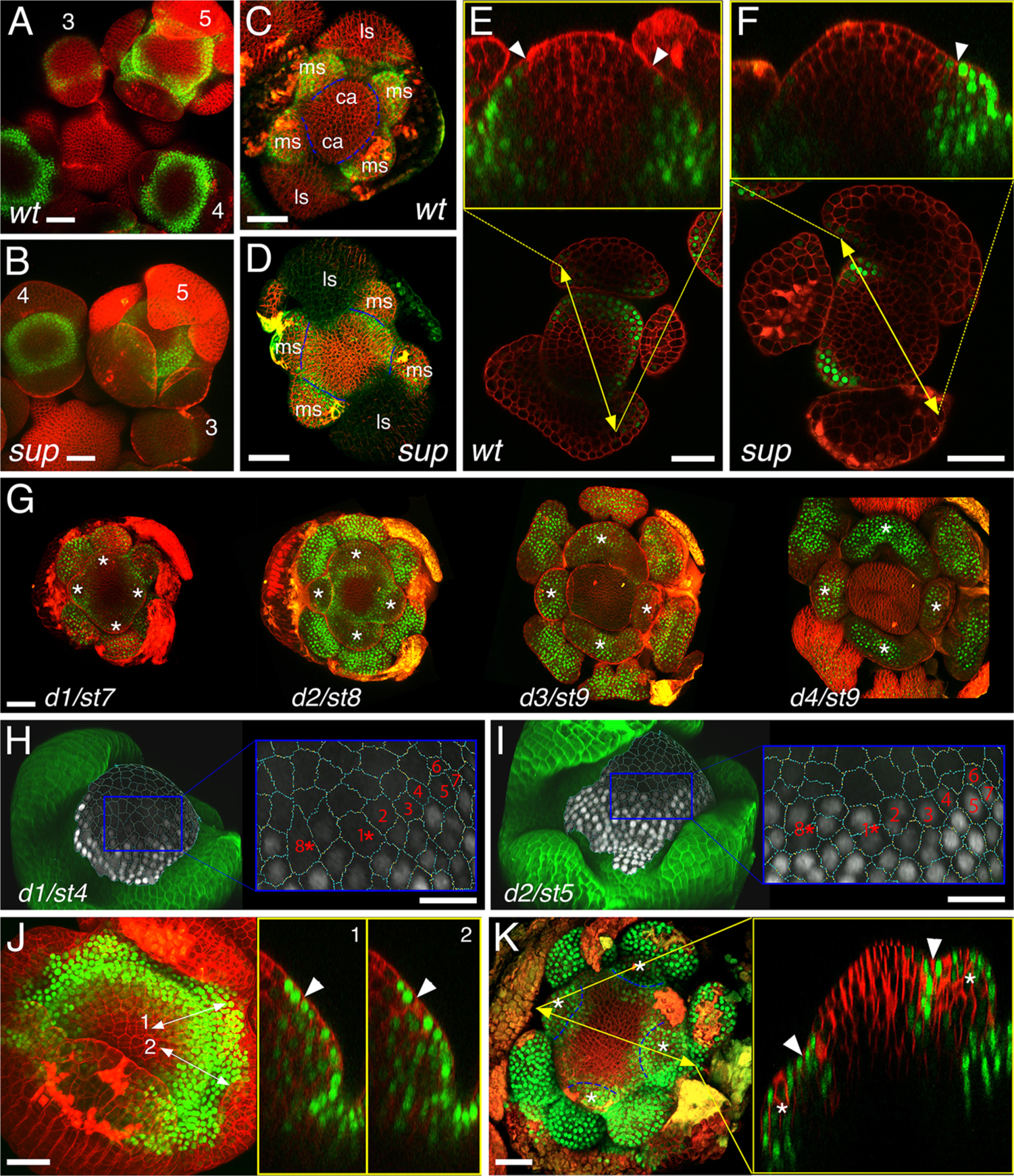
Expression of *AP3* in wild-type and *sup-1* flowers. Expression of the *pAP3-3xVenusN7* reporter in the wild type (A, C and E) and *sup-1* (B, D, F-K). (A-B) whole inflorescences; numbers indicate floral stages. (C-D) stage 5 flowers after removal of medial sepals; ca: carpel; ls: lateral sepal, covering lateral stamen; ms: medial stamen; dotted blue lines mark the boundary between whorl 3 stamens and the center of the flower. (E-F) stage 5 flowers, slice views along horizontal planes (bottom panels) and vertical planes (top panels); white arrowheads mark the boundary between whorl 3 stamens and the center of the flower. (G) 4-day time lapse of a single *sup-1* flower between stage 7 and stage 9; white asterisks mark extra stamen primordia. (H-I) 2-day time lapse of an individual *sup-1* flower between stage 4 (H) and 5 (I); left panels show a lateral view of the flower, with a segmented projection of the L1 layer in the center; right panels show a close-up of the same area on each day; red numbers mark cells that do not express *AP3* at stage 4, but express *AP3* at stage 5; asterisks indicate divisions that occurred between stage 4 and 5. (J) MaxIP (left panel) and slice view along vertical planes (right panels) of the flower shown in J; white arrowheads mark the boundary between whorl 3 stamens and the center of the flower. (K) MaxIP (left panel) and slice view along the vertical planes (right panel) of a stage 8 *sup-1* flower; white asterisks mark extra stamen primordia; white arrowheads and dashed blue lines mark the boundary between extra stamen primordia and the center of the flower. Bars = 20 µm.

We sought to establish whether the extra *AP3*-expressing cells in stage 5 *sup-1* flowers derive from whorl 4 cells that change identity, or from whorl 3 cells that overproliferate. The ring of extra *AP3*-expressing cells in *sup-1* flowers looks similar to the ring of *SUP*-expressing cells in whorl 4 of wild-type flowers (compare Fig. 2F to Fig. 1, D and I), suggesting that the loss of *SUP* function might cause ectopic expression of *AP3* in these cells. Using time-lapse imaging of *sup-1 pAP3-3xVenusN7* flower buds, we identified numerous individual cells at the boundary between whorls 3 and 4 that do not express *AP3* at stage 4 but begin to express *AP3 de novo* at stage 5 (Fig. 2, compare H and I). These cells that switch identity from female-fated, non *AP3*-expressing cells to male-fated, *AP3-*expressing cells are situated inside of the boundary between whorl 3 stamens and the center of the flower, indicating that they belong to whorl 4 (Fig. 2J). These data clearly show that the extra *AP3*-expressing cells in *sup-1* flowers originate from whorl 4 cells that switch fate from female to male, rather than from whorl 3 cells that overproliferate, and supports the whorl 4 model.

### Stem cell termination is delayed in sup-1 flowers

The respecification of a small ring of cells in the fourth whorl of *sup-1* flowers at stage 5 is not sufficient to explain the formation of so many supernumerary stamens. The iterative production of rings of extra stamens in the fourth whorl of *sup-1* flowers requires an increase or prolongation of cell proliferation compared to the wild type. To test if cells in the floral meristem are the source of overproliferation in *sup-1* mutants, we monitored the expression of stem cell marker *CLAVATA3* (*CLV3*) and stem cell-promoting gene *WUSCHEL* (*WUS*) using *pCLV3-erGFP*^15^ and *pWUS-erGFP* transcriptional reporters. *CLV3* expression persists in wild-type flowers through stage 6 (Fig. 3A)^16^, but is no longer detectable at stage 7, as stem cells are incorporated into developing carpels (Fig. 3B). Conversely, we observed *CLV3* expression as late as stage 10 in a small dome at the center of *sup-1* flowers, after several extra stamens have formed (Fig. 3C). Stem cell termination is thus clearly delayed in *sup-1* flowers compared to the wild type. Similarly, *WUS* expression stops by stage 5 in wild-type flowers^17^, but is maintained much longer in some *sup-1* flowers (Fig. 3D), indicating that a *bona fide* FM remains functional in *sup-1* flowers longer than it does in the wild type. *AG* is responsible for triggering stem cell termination in wild-type flowers by turning off the expression of *WUS*^18–21^, and most mutants with a delay or loss of floral stem cell termination have defects in *AG* expression^4^. We thus used the *gAG-GFP* reporter^22^ to compare the expression of *AG* in wild-type and *sup-1* flowers, and *AG* expression appears unaffected in *sup-1* flowers (Fig. 3, compare E and F and G and H)^10^, suggesting that an *AG*-independent mechanism is responsible for the delay in stem cell termination in *sup-1* flowers. *SUP* affects floral stem cells non cell-autonomously, as *SUP* and *CLV3* expression domains are separated by a narrow, 1-2-cell wide ring (Fig. 3I; Fig. S8A). Indeed, this ring of cells separating the *SUP* and *CLV3* expression domains expresses *DRNL* and likely corresponds to the carpel founder cells (Fig. S4, D-E). However, the *SUP* expression domain tightly surrounds *WUS* expression, with a few cells expressing both genes, suggesting that the effect of *SUP* on stem cells may be mediated by *WUS* (Fig. S8B; Movie S2).

**Figure 3.**
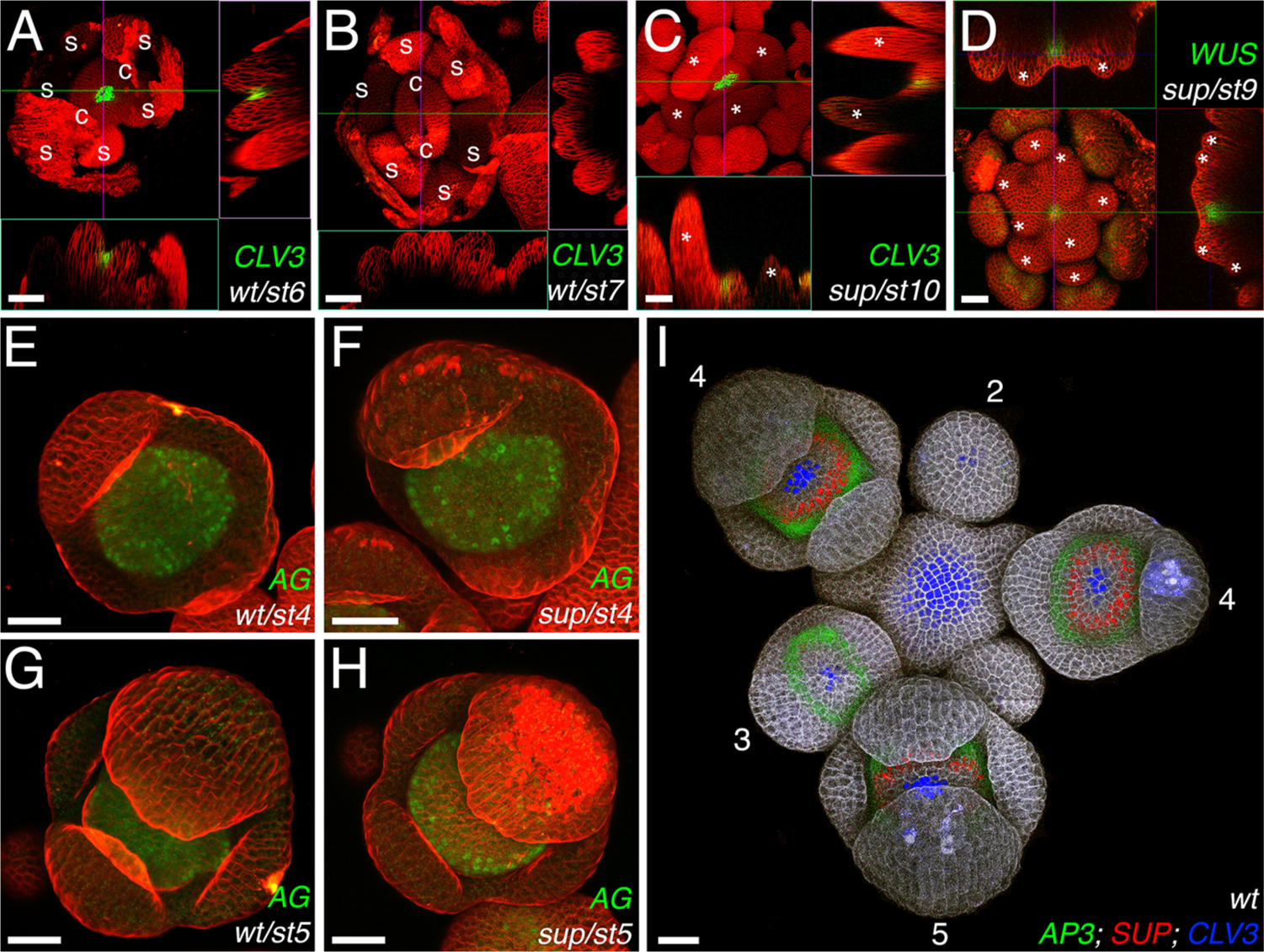
*SUP* promotes stem cell termination non-cell autonomously, and independently of *AG* expression. (A-C) expression of the *pCLV3-erGFP* reporter in stage 6 (A) and 7 (B) wild-type flowers, and in a stage 10 *sup-1* flower (C); top left panels show MaxIPs, with GFP fluorescence detected with Imaris; bottom left and top right panels show slice views along the xz and yz planes, respectively; s: whorl 3 stamen, c: carpel. (D) expression of the *pWUS-erGFP* reporter in a stage 8 *sup-1* flower. Asterisks in C and D mark extra stamen primordia. (E-H) expression of the *gAG-GFP* reporter in stage 4 (E and F) and 5 (G and H) wild-type (E-F) and *sup-1* (G-H) flowers. (I) expression of the *gAP3-GFP* (green), *gSUP-3xVenusN7* (red) and *pCLV3-dsRedN7* (blue)^30^ reporters in a wild type inflorescence; numbers indicate floral stages. Bars = 20 µm.

## Discussion

It is worth noting that several studies have shown that ectopic expression of *SUP* causes a decrease in cell proliferation^5,23–25^, which was interpreted as evidence in support of the whorl 3 model. It is not surprising, however, for a boundary gene to control cell proliferation, as cell division rates are lower at boundaries, including the boundary between stamens and carpels, than in developing organs^2,26^. For instance, *RABBIT EARS* (*RBE*), a gene closely related to *SUP*, specifies the boundary between whorls 2 and 3 by excluding *AG* from whorl 2^27,28^, and also specifies the intersepal boundaries by regulating cell proliferation in whorl 1 via the miR164/*CUP-SHAPED COTYLEDON* (*CUC*) module^29^. Similarly, a role for *SUP* in the control of cell proliferation does not exclude the possibility that *SUP* also affects *AP3/PI* expression. Moreover, rates of cell proliferation on either side of the boundary between whorls 3 and 4 appear unaffected in *sup-1* flowers compared to the wild type^26^, contrary to the predictions of the whorl 3 model.

Our data confirm, instead, the predictions of the whorl 4 model. Specifically, we show that the extra stamens in *sup* mutant flowers arise from a narrow ring of cells in the outer part of whorl 4, adjacent to the boundary with whorl 3, which change identity from female to male at the transition between stages 4 and 5, and start expressing *AP3 de novo* (Fig. 2). Cells in this ring then divide, allowing for the formation of extra stamens. The *sup* phenotype was initially described as heterochronic, *sup* flowers being “stuck in developmental time”^10,11^. The *sup* phenotype is indeed iterative: as extra stamen primordia arise, the lack of functional *SUP* at the inner boundary of these stamens causes *AP3* expression to spread again toward the center of the flower (Fig. 2K), allowing for the formation of additional extra stamens. Even as several rings of extra stamens form one after the other, the center of the flower, which is still devoid of *AP3* expression, is replenished by the floral stem cells, which are maintained longer in *sup* flowers than in the wild type (Fig. 3). Eventually, the center of *sup* flowers differentiates into stunted carpels or mosaic, stamen-carpel organs (Fig. S7). The fact that *SUP* is expressed in the fourth whorl of wild-type flowers, in the same cells that express *AP3* in the fourth whorl of *sup* flowers (Fig. 1, D and I; Fig. 2, D and F), suggests that *SUP* cell-autonomously represses *AP3* expression in the outer part of whorl 4. Whether such a repression is direct or indirect, however, remains unknown. Conversely, *SUP* affects floral stem cells non cell-autonomously, and independently of *AG* expression (Fig. 3).

Over the last two decades, tremendous effort has been put into understanding the mechanisms that underlie the formation of boundaries between different organs and between organs and the meristem, both in the SAM and the FM ^2,4^. Numerous genes have been characterized, with some, like the *CUC* genes, involved in the formation of all boundaries, and some, like *SUP* or *RBE*, involved in the formation of specific boundaries in the flower. However, most of these genes have been associated with the growth suppression aspect, and not the identity separation aspect of boundary formation ^2,4^. This study provides new insight into how a boundary gene separates two different developmental programs in adjacent organs.

## Acknowledgments

The authors wish to thank John Chandler and Paul Tarr for providing reporter lines, Richard Immink for providing the unpublished *gAP3-GFP* and *gPI-GFP* reporter lines, Toshiro Ito and Frank Wellmer for critical reading of the manuscript, and Ann Lavanway, a.k.a. Dartmouth Biology Department’s Superwoman, for her tremendous help with live confocal imaging. Funding in the Jack lab was provided by the US National Science Foundation through grant IOS-0926347. Funding in the Meyerowitz Laboratory was provided by the Howard Hughes Medical Institute, the US National Institutes of Health through grant R01 GM104244 and the Gordon and Betty Moore Foundation through Grant GBMF3406.

## Author contributions

NP, WY and TPJ performed the experiments (NP and TPJ generated the reporter lines; NP did the live confocal imaging and WY did the *in situ* hybridizations); NP, PD and TPJ designed the experiments; NP wrote the manuscript; PD, EMM and TPJ edited the manuscript.

## Data availability

The authors declare that all the data supporting the findings of this study are available within the paper and its supplementary information files.

## Supplemental Materials

Methods

Figures S1-S8

Table S1

Movies S1-S2

## References

1. Fagotto, F. The cellular basis of tissue separation. Development 141, 3303–3318 (2014).

2. Aida, M. & Tasaka, M. Morphogenesis and patterning at the organ boundaries in the higher plant shoot apex. Plant Mol. Biol. 60, 915–928 (2006).

3. Krizek, B. A. & Fletcher, J. C. Molecular mechanisms of flower development: an armchair guide. Nat. Rev. Genet. 6, 688–698 (2005).

4. Prunet, N. & Jack, T. P. Flower Development in Arabidopsis: There is more to it than learning your ABCs. Methods Mol. Biol. 1110, 3–33 (2014).

5. Hiratsu, K., Ohta, M., Matsui, K. & Ohme-Takagi, M. The SUPERMAN protein is an active repressor whose carboxy-terminal repression domain is required for the development of normal flowers. FEBS Letters 514, 351–354 (2002).

6. Sakai, H., Medrano, L. J. & Meyerowitz, E. M. Role of SUPERMAN in maintaining Arabidopsis floral whorl boundaries. Nature 378, 199–203 (1995).

7. Dathan, N. et al. The Arabidopsis SUPERMAN protein is able to specifically bind DNA through its single Cys2-His2 zinc finger motif. Nuc. Ac. Res. 30, 4945–4951 (2002).

8. Hiratsu, K., Mitsuda, N., Matsui, K. & Ohme-Takagi, M. Identification of the minimal repression domain of SUPERMAN shows that the DLELRL hexapeptide is both necessary and sufficient for repression of transcription in Arabidopsis. Biochem. Biophys. Res. Com. 321, 172–178 (2004).

9. Ito, T., Sakai, H. & Meyerowitz, E. M. Whorl-specific expression of the SUPERMAN gene of Arabidopsis is mediated by cis elements in the transcribed region. Curr. Biol. 13, 1524–1530 (2003).

10. Bowman, J. L. et al. SUPERMAN, a regulator of floral homeotic genes in Arabidopsis. Development 114, 599–615 (1992).

11. Schultz, E. A., Pickett, F. B. & Haughn, G. W. The FLO10 Gene Product Regulates the Expression Domain of Homeotic Genes AP3 and PI in Arabidopsis Flowers. Plant Cell 3, 1221–1237 (1991).

12. Sakai, H., Krizek, B. A., Jacobsen, S. E. & Meyerowitz, E. M. Regulation of SUP expression identifies multiple regulators involved in arabidopsis floral meristem development. Plant Cell 12, 1607–1618 (2000).

13. Smyth, D. R., Bowman, J. L. & Meyerowitz, E. M. Early flower development in Arabidopsis. Plant Cell 2, 755–767 (1990).

14. Chandler, J. W., Jacobs, B., Cole, M., Comelli, P. & Werr, W. DORNROSCHEN-LIKE expression marks Arabidopsis floral organ founder cells and precedes auxin response maxima. Plant Mol. Biol. 76, 171–185 (2011).

15. Reddy, G. V. & Meyerowitz, E. M. Stem-cell homeostasis and growth dynamics can be uncoupled in the Arabidopsis shoot apex. Science 310, 663–667 (2005).

16. Fletcher, J. C., Brand, U., Running, M. P., Simon, R. & Meyerowitz, E. M. Signaling of cell fate decisions by CLAVATA3 in Arabidopsis shoot meristems. Science 283, 1911–1914 (1999).

17. Mayer, K. F. et al. Role of WUSCHEL in regulating stem cell fate in the Arabidopsis shoot meristem. Cell 95, 805–815 (1998).

18. Lenhard, M., Bohnert, A., Jurgens, G. & Laux, T. Termination of stem cell maintenance in Arabidopsis floral meristems by interactions between WUSCHEL and AGAMOUS. Cell 105, 805–814 (2001).

19. Lohmann, J. U. et al. A molecular link between stem cell regulation and floral patterning in Arabidopsis. Cell 105, 793–803 (2001).

20. Liu, X. et al. AGAMOUS terminates floral stem cell maintenance in Arabidopsis by directly repressing WUSCHEL through recruitment of Polycomb Group proteins. Plant Cell 23, 3654–3670 (2011).

21. Sun, B., Xu, Y., Ng, K. H. & Ito, T. A timing mechanism for stem cell maintenance and differentiation in the Arabidopsis floral meristem. Genes & Dev. 23, 1791–1804 (2009).

22. Urbanus, S. L. et al. In planta localisation patterns of MADS domain proteins during floral development in Arabidopsis thaliana. BMC Plant Biol. 9, 5 (2009).

23. Bereterbide, A., Hernould, M., Castera, S. & Mouras, A. Inhibition of cell proliferation, cell expansion and differentiation by the Arabidopsis SUPERMAN gene in transgenic tobacco plants. Planta 214, 22–29 (2001).

24. Nandi, A. K., Kushalappa, K., Prasad, K. & Vijayraghavan, U. A conserved function for Arabidopsis SUPERMAN in regulating floral-whorl cell proliferation in rice, a monocotyledonous plant. Curr. Biol. 10, 215–218 (2000).

25. Yun, J. Y., Weigel, D. & Lee, I. Ectopic expression of SUPERMAN suppresses development of petals and stamens. Plant & Cell Physiol. 43, 52–57 (2002).

26. Breuil-Broyer, S. et al. High-resolution boundary analysis during Arabidopsis thaliana flower development. Plant J. 38, 182–192 (2004).

27. Krizek, B. A., Lewis, M. W. & Fletcher, J. C. RABBIT EARS is a second-whorl repressor of AGAMOUS that maintains spatial boundaries in Arabidopsis flowers. Plant J. 45, 369–383 (2006).

28. Takeda, S., Matsumoto, N. & Okada, K. RABBIT EARS, encoding a SUPERMAN-like zinc finger protein, regulates petal development in Arabidopsis thaliana. Development 131, 425–434 (2004).

29. Huang, T., Lopez-Giraldez, F., Townsend, J. P. & Irish, V. F. RBE controls microRNA164 expression to effect floral organogenesis. Development 139, 2161–2169 (2012).

30. Zhou, Y. et al. Control of plant stem cell function by conserved interacting transcriptional regulators. Nature 517, 377–380 (2015).

